# ppGpp is Present in and Functions to Regulate Sleep in *Drosophila*

**DOI:** 10.1101/2022.11.17.516975

**Authors:** Xihuimin Dai, Wei Yang, Xiaohui Zhang, Enxing Zhou, Ying Liu, Tao Wang, Wenxia Zhang, Xinxiang Zhang, Yi Rao

## Abstract

Sleep is essential for animals, and receives inputs from circadian, homeostasis, and environment, yet the mechanisms of sleep regulation remain elusive. Discovery of molecules in living systems and demonstration of their functional roles are pivotal in furthering our understanding of the molecular basis of biology. Here we report that ppGpp (guanosine-5’-diphosphate, 3’-diphosphate), a molecule that has been detected in prokaryotes for more than five decades, is present in *Drosophila*, and plays an important role in regulation of sleep and SISL (starvation induced sleep loss). ppGpp is detected in germ-free *Drosophila* and hydrolyzed by an enzyme encoded by the *mesh1* gene in *Drosophila*. Nighttime sleep and SISL were defected in *mesh1* mutant flies, and rescued by expression of wildtype Mesh1, but not the enzymatically defective mutant Mesh1E66A. Ectopic expression of RelA, the *E. coli* synthetase for ppGpp, phenocopied *mesh1* knockout mutants, whereas overexpression of Mesh1 resulted in the opposite phenotypes, supporting that ppGpp is both necessary and sufficient in sleep regulation. A chemoconnectomic screen followed by genetic intersection experiments implicate the Dilp2 neurons in the *pars intercerebralis* (PI) brain region as the site of ppGpp function. Our results have thus supported that ppGpp is present in animals after long lag since its discovery in bacteria, and revealed a physiological role of ppGpp in sleep regulation for the first time.

## INTRODUCTION

It was 50 years ago when guanosine-5’-diphosphate, 3’-diphosphate (guanosine tetraphosphate, ppGpp) and guanosine-5’-triphosphate, 3’-diphosphate (guanosine pentaphosphate, pppGpp) were implicated in gene regulation in *Escherichia coli* (*E. coli*)(Cashel and Gallant, 1969). Collectively known as (p)ppGpp, they are key players in bacterial stringent response to amino acid starvation(Dalebroux and Swanson, 2012; Field, 2018; Gourse et al., 2018; Hauryliuk et al., 2015; Liu et al., 2015; Magnusson et al., 2005; Potrykus and Cashel, 2008; Wang et al., 2007). The level of ppGpp is regulated by the RelA/SpoT Homolog (RSH) family(Haseltine and Block, 1973; Hogg et al., 2004). In bacteria, RelA is one of the RSHs(Haseltine and Block, 1973), which contains both a ppGpp synthetase domain (SYNTH) and an inactive ppGpp hydrolase domain (HD)(Hogg et al., 2004). When amino acids are depleted, uncharged tRNAs accumulate(Fangman and Neidhardt, 1964) and activate the ribosome-associated RelA(Yang and Ishiguro, 2001), which increases the production of ppGpp(Cochran and Byrne, 1974). Another typical RSH in bacteria is the ppGpp hydrolase SpoT(Laffler and Gallant, 1974), containing a weak SYNTH(Leung and Yamazaki, 1977) and an active HD(Murray and Bremer, 1996). Upon iron limitation(Vinella et al., 2005), carbon starvation(Lesley and Shapiro, 2008), or glucose phosphate stress(Kessler et al., 2017), ppGpp was also increased(Potrykus and Cashel, 2008).

ppGpp has also been detected in plants(Boniecka et al., 2017; Ito et al., 2012; Kasai et al., 2004; Sugliani et al., 2016; Tozawa et al., 2007; van der Biezen et al., 2000; Xiong et al., 2001). More than 30 members of the RSHs have been found in bacteria and plants(Atkinson et al., 2011; Field, 2018).

(p)ppGpp has not been detected in animals until very recently. Early detection of ppGpp in the mouse turned out to be irreproducible(Irr et al., 1974; Martini et al., 1977; Silverman and Atherly, 1977). Work in mammalian cell lines also failed to detect ppGpp before or after amino acid deprivation(Dabrowska et al., 2006; Fan et al., 1973; Givens et al., 2004; Kim et al., 2009; Mamont et al., 1972; Rapaport et al., 1975; Sato et al., 2015; Smulson, 1970; Thammana et al., 1976; Thompson et al., 1973; Yamada et al., 2003). In *Drosophila*, metazoan SpoT homolog-1 (Mesh1), a member of the RSH family containing only the hydrolase domain for ppGpp has been found(Sun et al., 2010). And during the submission of this work, ppGpp has just been reported to be detected in *Drosophila* and human cells, and plays an important role in metabolism (Ito et al., 2020).

*Drosophila* has been served as a model for genetic studies of sleep for more than two decades(Hendricks et al., 2000; Shaw and Brody, 2000). Fly sleep is regulated by multiple genes functioning in several brain regions such as the fan-shape bodies (FSBs), the ellipsoid body (EB), the mushroom bodies (MB), and the *pars intercerebralis* (PI) (Artiushin and Sehgal, 2017; Shafer and Keene, 2021). Beside the internal control from circadian and homeostasis, sleep is also regulated by environmental factors such as food. Starvation has been found to induce sleep loss in both flies and mammals(Danguir and Nicolaidis, 1979; Keene et al., 2010).

We have carried out a genetic screen for genes involved in sleep regulation(Dai et al., 2021; Dai et al., 2019) and here report that ppGpp is present in flies and regulates both daily sleep and starvation induced sleep loss. *mesh1*, the gene encoding ppGpp hydrolase, is expressed in a specific population of neurons, and the effects of RelA, the bacterial ppGpp synthetase, and Mesh1 overexpression could be detected when they were expressed in neurons, but not in non-neuronal cells. Further dissection narrowed down the functional significance of *mesh1*-expressing neurons in the PI. Thus, after more than 50 years of its discovery, we have confirmed the presence of ppGpp in animals and first found that it functions in sleep regulation in specific neurons.

## RESULTS

### ppGpp is present and regulated by *mesh1* in *Drosophila*

We screened through 1765 P-element insertion lines of *Drosophila* (Eddison et al., 2012) for mutations affecting sleep latency (Fig. 1A), and found that an insertion in the *mesh1* gene (*mesh1*-ins) (Fig. 1C) resulted in significantly longer sleep latency (Figs. 1A and D) and less total sleep duration (Fig. 1B).

**Figure 1.**
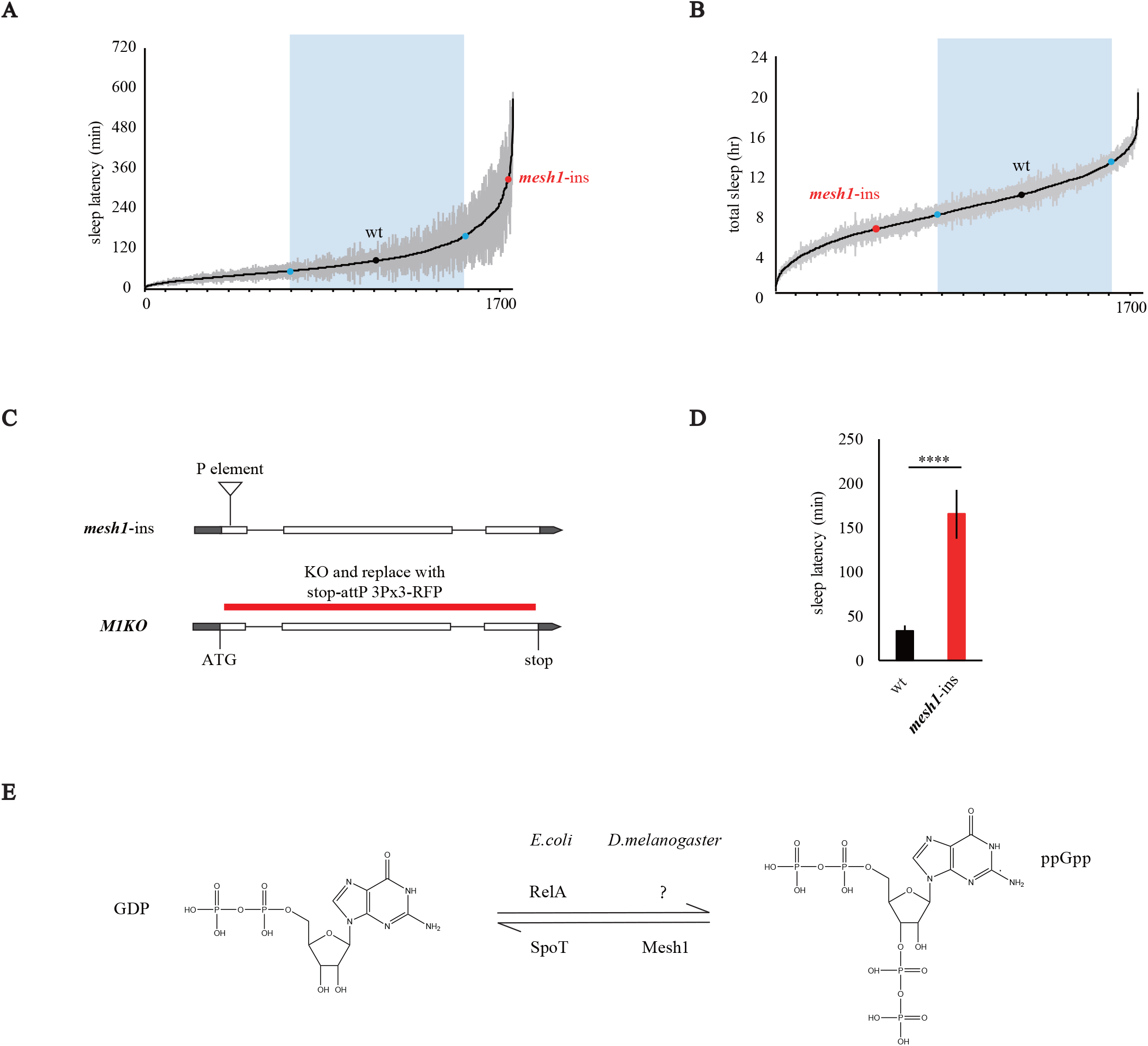
Results of a sleep latency screen. **(A-B)** Sleep latency **(A)** and total sleep duration **(B)** results of a screen of 1765 P-element insertion lines. *mesh1-ins* flies had significant longer sleep latency **(A)** and less total sleep **(B)** than wt flies. y axis is sleep latency in minutes and total sleep in hours respectively, and x axis is screened P element insertion lines. Grey shadows indicate the standard error of the mean (SEM) from multiple flies of each line, and blue dots and rectangles shows the range of 3-fold of standard deviation from the value of wt. **(C)** Schematic representations of *mesh1-*ins and *M1KO* genotypes. In *mesh1-*ins, the P-element was inserted into the CDS of the first exon. In *M1KO*, the entire CDS except the start codon was replaced with stop-2A-attP and 3Px3-RFP. **(D)** Statistical analysis. Sleep latency was significantly longer in *mesh1-*ins flies (Student’s t-test, **** *P* < 0.0001). Error bars represent s.e.m. **(E)** A diagram of ppGpp metabolism. In *E. coli*, GDP is converted to ppGpp by its synthetase RelA, whereas ppGpp is converted to GDP by the hydrolase SpoT. In *D. melanogaster*, only the hydrolase Mesh1 has been discovered but the synthetase is unknown.

*mesh1* encodes an RSH family member in animals, and Mesh1 protein is predicted to have hydrolase activity, converting ppGpp to GDP(Sun et al., 2010)(Fig. 1E). However, ppGpp has not been detected in animals until very recently (Ito et al., 2020), and the function of Mesh1 remains largely unknown..

Through expression, purification, and an *in vitro* hydrolysis assay of *Drosophila* Mesh1 protein in *E. coli*, we found that ppGpp was indeed degraded by Mesh1 *in vitro* (Fig. S1A). To test the existence of endogenous ppGpp in *Drosophila*, extracts from more than 2000 flies were made. ppGpp was detected by ultra-performance liquid chromatography and mass spectrometry (UPLC-MS) in wild type (wt) flies (Figs. 2A and B). We generated a knockout (KO) line of *mesh1* (*M1KO*) using CRISPR-Cas9 to replace most of its coding sequence following the start codon with 2A-attP (Fig. 1C), and found that more ppGpp was present in *M1KO* flies than the wt flies (Fig. 2C), consistent with the fact that *Drosophila mesh1* encodes only a hydrolase domain (Fig. S1B). Thus, the ppGpp detected by us was regulated by the *Drosophila mesh1* gene.

**Figure 2.**
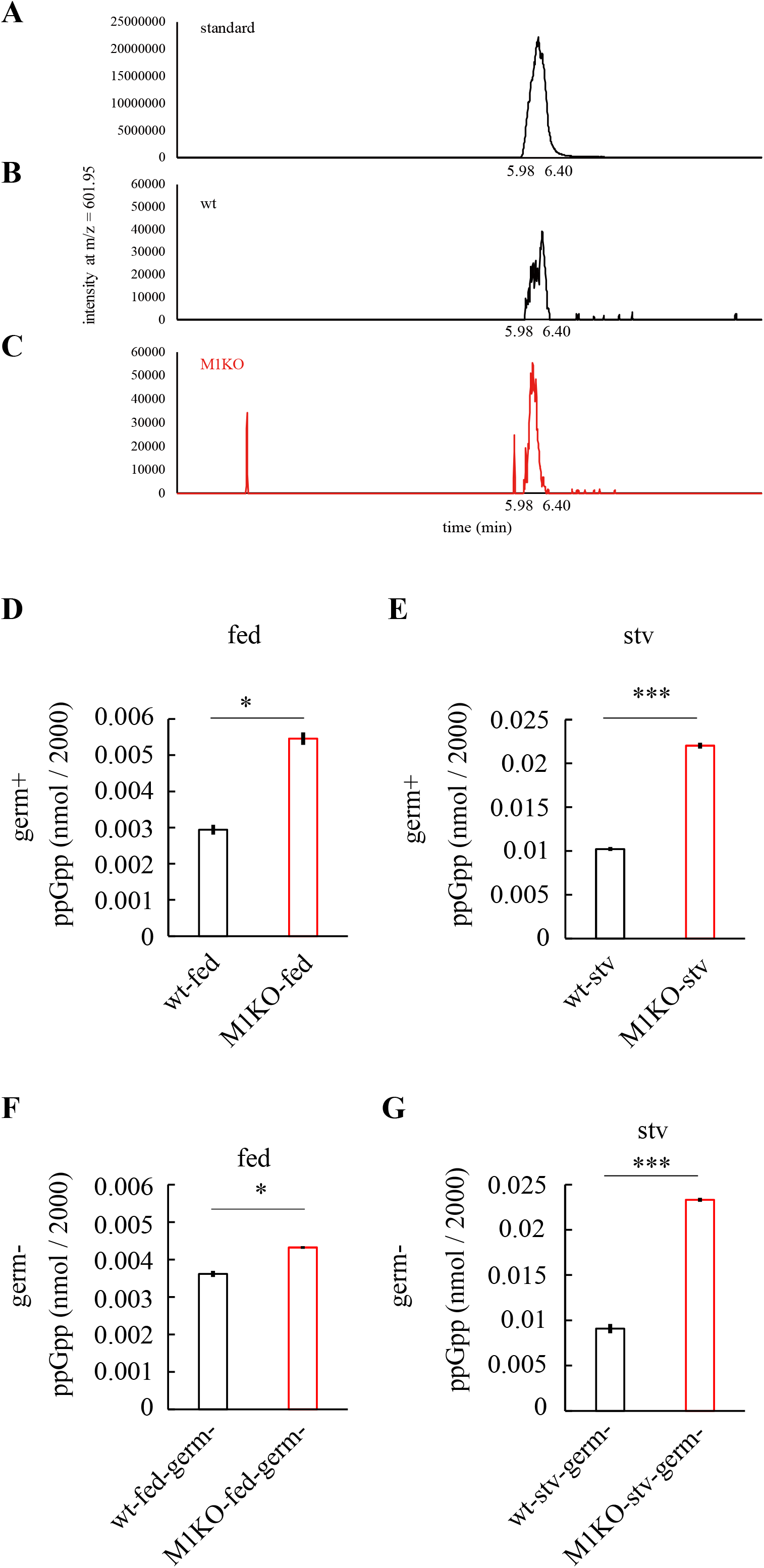
Identification and quantitative analysis of ppGpp in *Drosophila*. **(A-C)** UPLC-MS profiles of standard ppGpp **(A)**, extracts of wt flies**(B)**, and extracts of *M1KO* flies **(C)**. ppGpp is detected in wt flies and increased in M1KO flies. x axis is time (minutes), and y axis is intensity of MS signal at m/z = 601.95, which corresponds to standard ppGpp **(A)**. **(D-G)** Statistical analyses of ppGpp level under fed **(D**,**F)** and starved **(E**,**G)** conditions in normal flies **(D**,**E)** and germ-free flies **(F**,**G)**. For each group, ∼2000 male flies were sacrificed for UPLC-MS measurements. In all conditions, ppGpp level was significantly increased in *M1KO* flies compared with wt flies. Student’s t-test was used, * *P* < 0.05, *** *P* < 0.001. Error bars represent s.e.m.

To tell whether the detected ppGpp was synthesized by the flies or from bacteria attached to the flies, we generated germ-free lines of wt and *M1KO* for analysis. We confirmed that the flies were indeed germ-free flies with both bacterial culture (Fig. S1C) and 16S rDNA PCR (Fig. S1D). Because ppGpp is known to be involved in starvation response in bacteria(Potrykus and Cashel, 2008; Ronneau and Hallez, 2019), and *mesh1* mutant flies have been reported to have impaired starvation resistance(Sun et al., 2010), we tested ppGpp in fed and starved flies. ppGpp was detected in both fed and starved germ-free flies, and levels in *M1KO* were significantly higher in both conditions (Figs. 2D-G), suggesting that flies can indeed synthesize ppGpp.

Taken together, our results indicate that ppGpp is present in *Drosophila* and is regulated by *mesh1* gene.

### Decreased sleep at early night and late night in *mesh1* mutants

Because we identified mesh1 mutant through a sleep screen, we then looked at the sleep profiles of *M1KO* mutant flies. *M1KO* flies sleep significantly less than wt flies, and showed shifted activity peaks (Fig. 3A). In the paradigm of 12 hours light and 12 hours darkness (LD), *M1KO* flies showed significantly decreased nighttime sleep (Fig. 3B), decreased total sleep (Fig. 3D) and increased sleep latency at night (Fig. 3E), but no significant change in daytime sleep level (Fig. 3C) or daytime sleep latency (Fig. 3F) was detected. Sleep bout number and length were not significantly different between wt and *M1KO* flies (Fig. S2).

**Figure 3.**
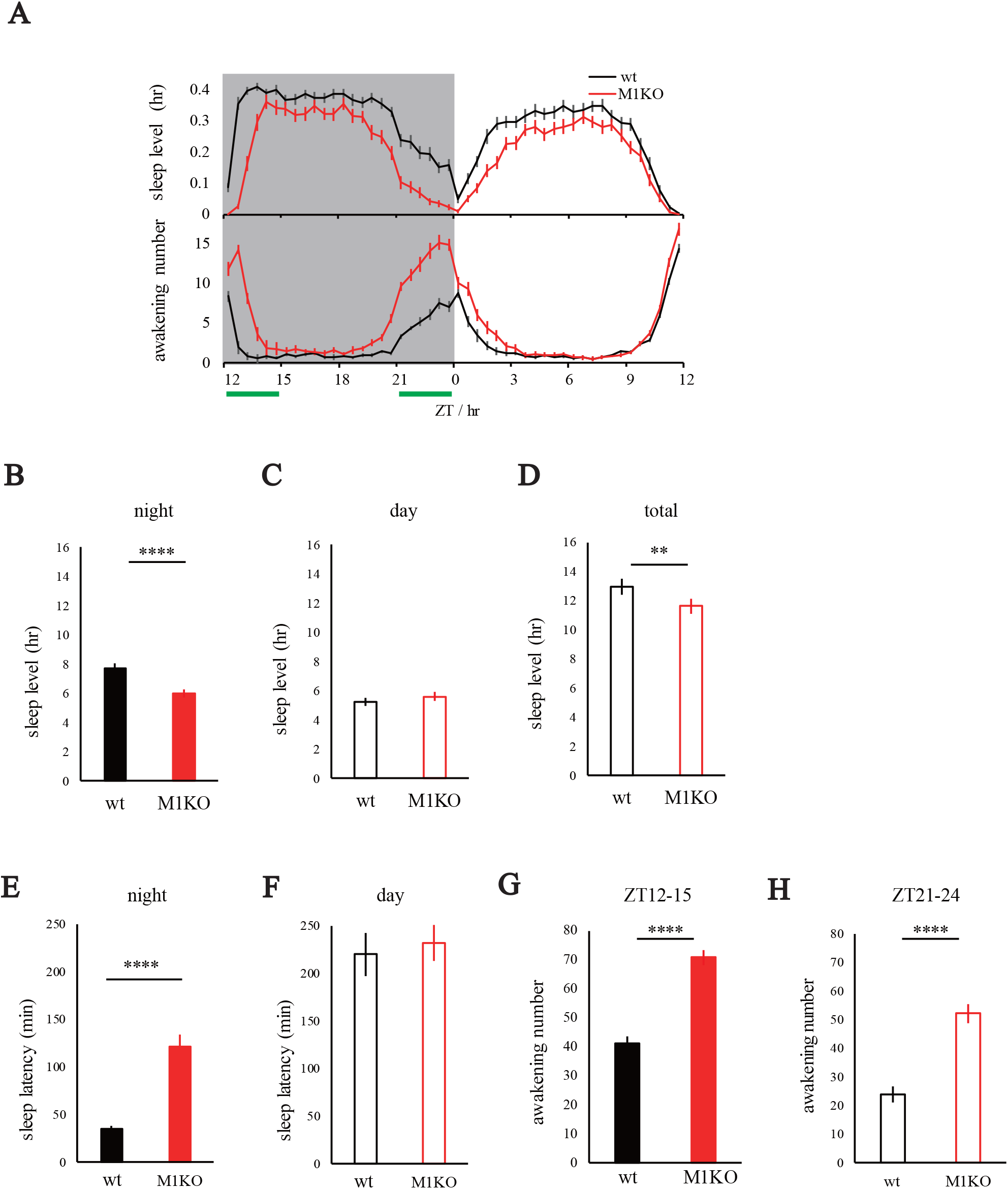
Sleep phenotypes of *M1KO* mutant flies. **(A)** Profiles of sleep (top) and awakening (bottom) in *M1KO* Flies, plotted in 30mins bins. Shaded background indicates the dark phase (ZT12-24); white background indicates the light phase (ZT0-12). Early night (ZT 12-15) and late night (ZT 21-24) were denoted with green bars. **(B-D)** Statistical analyses of sleep duration during nighttime **(B)**, daytime **(C)** and in total **(D)**. Nighttime and total sleep were significantly reduced in *M1KO* flies. **(E-F)** Statistical analyses of sleep latency of nighttime **(E)** and daytime **(F)**. Nighttime sleep latency was significantly increased in *M1KO* flies. **(G-H)** Statistical analyses of summed awakening numbers at early night **(G)** and late night **(H)**. Awakening numbers of both early night and late night were significantly increased in *M1KO* flies. Student’s t-test, ** *P* < 0.01, **** *P* < 0.0001. Error bars represent s.e.m.

We assessed awakening according to a procedure reported recently(Tabuchi et al., 2018). Awakening number at the beginning of nighttime sleep (early night) (Fig. 3G) and awakening number near the end of nighttime sleep (late night) (Fig. 3H) were significantly increased in *M1KO* mutants.

Taken together, these results showed that it takes longer for *M1KO* flies to fall sleep at early night, and they wake up earlier in the late night, indicating a role of *mesh1* in regulating sleep latency at early night and awakening at late night.

### The enzymatic activity of *mesh1* is essential to sleep

Next, we asked whether the enzymatic activity of *mesh1*, which hydrolyzes ppGpp, is important to sleep. A previous *in vitro* study has shown that an E66A mutation in Mesh1 protein significantly disruptes the ppGpp hydrolase activity(Sun et al., 2010). Consistent with the previous study, we found that when expressed in bacteria, Mesh1E66A could not hydrolyze ppGpp (Fig. S1A).

To enable manipulation of *mesh1*-expressing cells, we generated a KO-Gal4 line of *mesh1* (*M1KOGal4*) by replacing its CDS after the start codon with an in-frame fusion of the 2A peptide and the yeast transcription factor Gal4 (Fig. 4A). We then expressed wildtype Mesh1 and Mesh1E66A respectively in *mesh1*-expressing cells of the mutant flies driven by *M1KOGal4*. We found that compared with *M1KO* mutant flies (Fig. S1E, column 2), the level of ppGpp in flies was rescued to control group level (Fig. S1E, column 3) by expression of wildtype Mesh1 (Fig. S1E, column 4), but could not be rescued by expression of Mesh1E66A (Fig. S1E, column 6). These results indicate that wt Mesh1 protein could, but Mesh1 E66A mutant protein could not, hydrolyze ppGpp either *in vitro* or *in vivo*.

**Figure 4.**
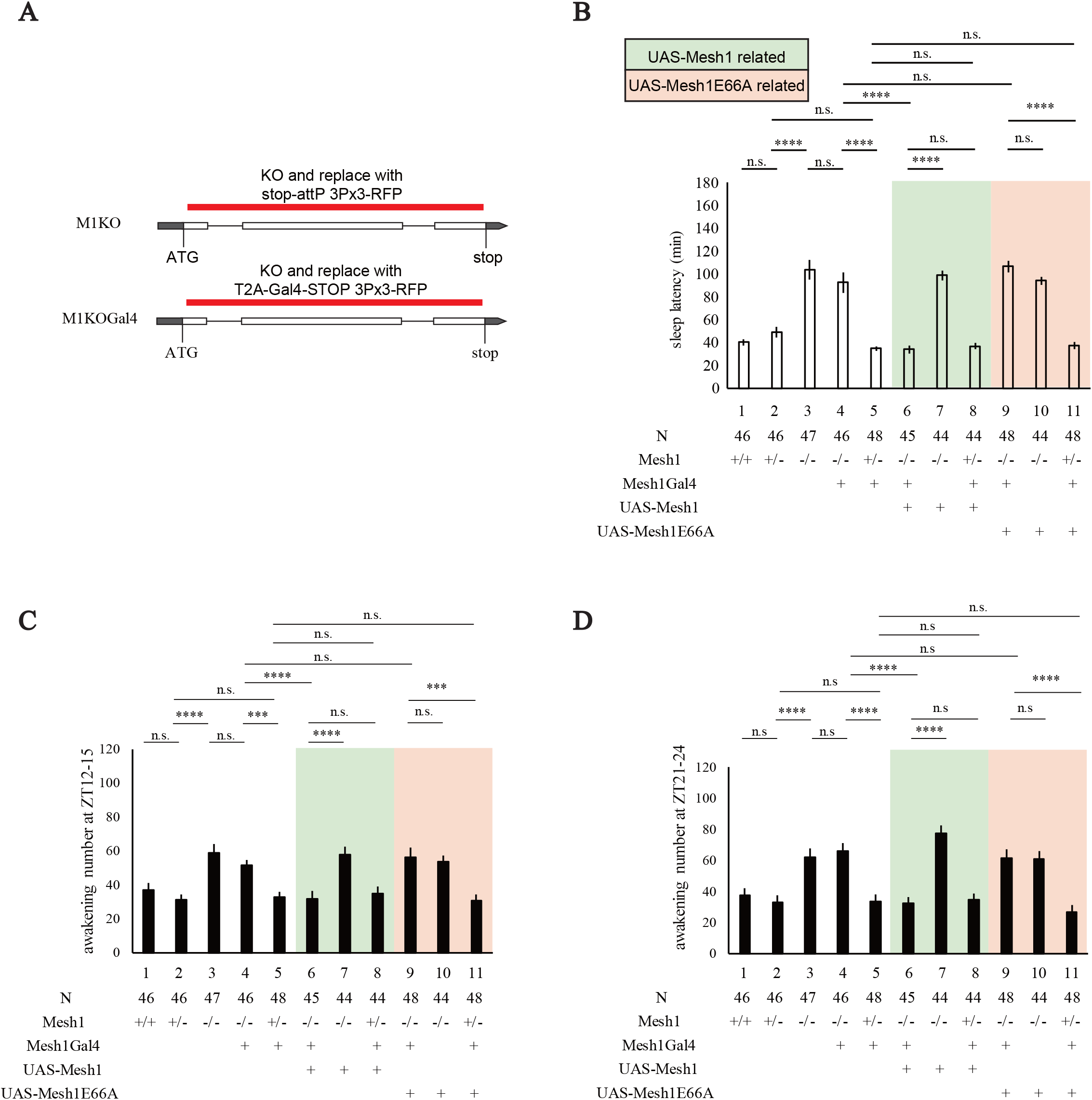
Significance of the hydrolysis activity of Mesh1 in sleep. **(A)** Schematic representations of *mesh1* gene with the red bar indicating the region deleted in *M1KO* and *M1KOGal4* flies respectively. *mesh1* CDS except the start codon was deleted in both *M1KO* and *M1KOGal4*, and the only difference is that a T2A-fused Gal4 (with a stop codon) were placed in *M1KOGal4*. **(B-D)** Sleep latency at night **(B)**, awakening number at early night **(C)**, and awakening number at late night **(D)** were rescued to wildtype level by expression of Mesh1 (column 6), but not Mesh1E66A (column 9). Numbers of flies used were denoted below each bar. Genotypes from left to right: **(1)** wildtype **(2)** *M1KO/+* **(3)** *M1KO/M1KO* **(4)** *M1KOGal4/M1KOGal4* **(5)** *M1KOGal4/+* **(6)** *M1KOGal4/M1KO, UAS-Mesh1* **(7)** *M1KO/M1KO, UAS-Mesh1* **(8)** *M1KOGal4/UAS-Mesh1* **(9)** *M1KOGal4/M1KO, UAS-Mesh1E66A* **(10)** *M1KO/M1KO, UAS-Mesh1E66A* **(11)** *M1KOGal4/UAS-Mesh1E66A*. Two-way ANOVA and Bonferroni post-tests was used, *** *P* < 0.001, **** *P* < 0.0001. Error bars represent s.e.m.

We then looked at whether wildtype Mesh1 and Mesh1E66A could rescue the sleep phenotypes of longer sleep latency (Fig. 4B), increased awakening number at early night (Fig. 4C) and late night (Fig. 4D). We found that expression of wildtype Mesh1 driven by *M1KOGal4* rescued all these phenotypes (Figs. 4B-D, column 6). By contrast, expression of Mesh1E66A could not rescue the sleep phenotypes in *M1KOGal4/M1KO* flies (Figs. 4B-D, column 9).

Taken together, the *in vitro* results from bacterially expressed Mesh1 and Mesh1E66A proteins and the *in vivo* results from genetic rescue experiments in flies strongly support that Mesh1 functions through ppGpp to regulate sleep.

### *mesh1* is expressed in the nervous system of *Drosophila*

To examine the expression pattern of Mesh1, we crossed *M1KOGal4* with each of the following four UAS lines: *UAS-mCD8-GFP* for membrane labeling(Lee and Luo, 1999), *UAS-redStinger* for nuclei labeling(Barolo et al., 2004), *UAS-denmark* for dendrites labeling(Nicolai et al., 2010) and *UAS-syt::eGFP* for axon terminals labeling(Zhang et al., 2002).

We found that Mesh1 is expressed in neurons in the central brain and the ventral nerve cord (Figs. 5A, B). In the central brain, *mesh1*-expressing neurons were detected in the PI and the suboesophageal ganglia (SOG) (Fig. 5E). The *mesh1*-expressing neurons in the PI (Fig. 5A) with their axonal terminals in the SOG (Fig. 5D, E) were reminiscent of the insulin-producing cells (IPC) in the PI which were shown previously to regulate sleep(Crocker et al., 2010).

**Figure 5.**
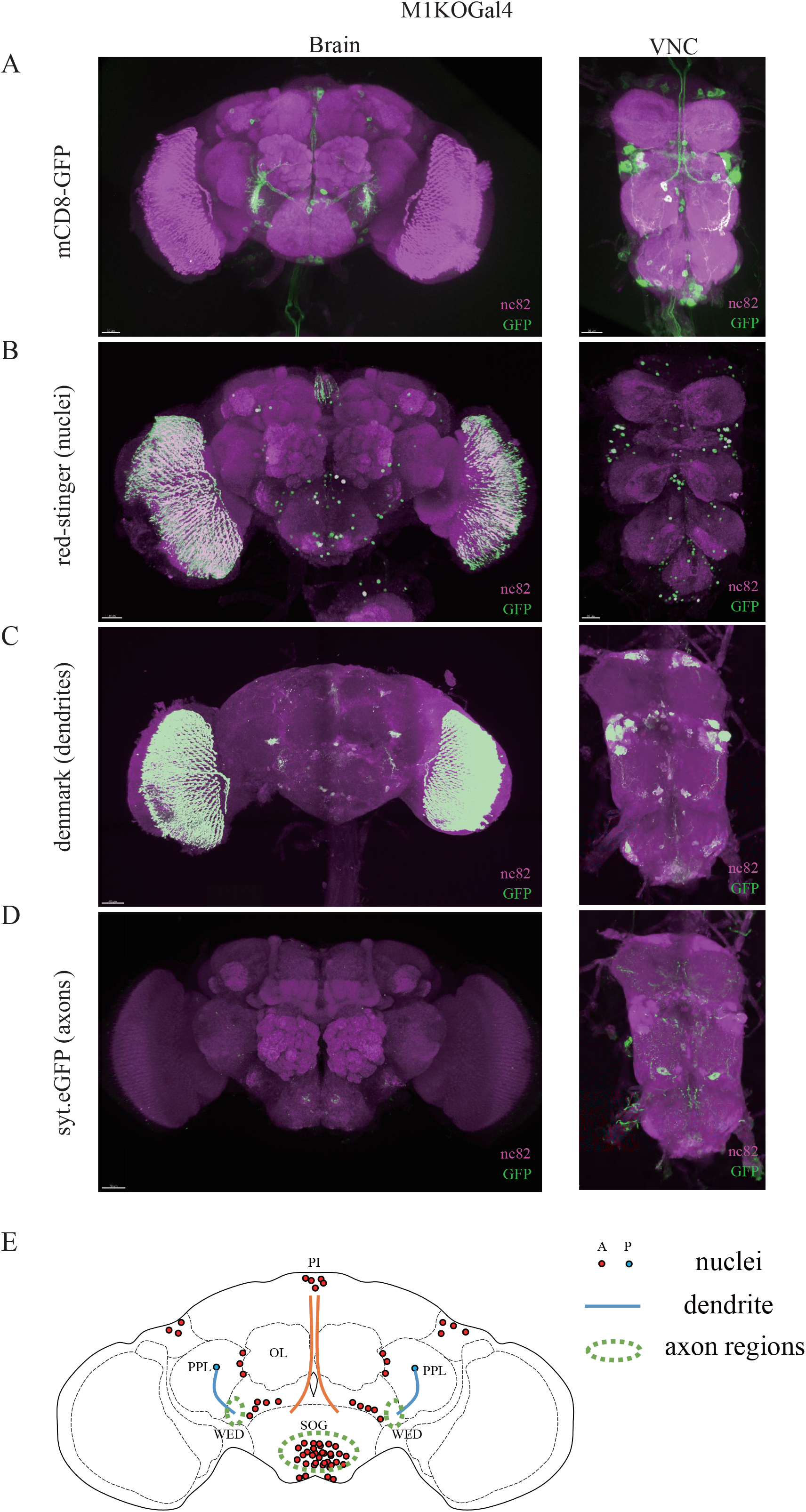
Expression patterns of *mesh1*. **(A-D)** Expression patterns of *M1KOGal4* labeled by mCD8-GFP **(A)**, red-stinger **(B)**, Denmark **(C)**, and Syt-eGFP **(D)** in the brain (left) and the VNC (right), immunostained with anti-GFP (green) and nc82 (magenta). **(E)** A diagrammatic summary of neurons labeled by *M1KOGal4*. A: anterior sections. P: posterior sections.

### Neuronal ppGpp regulates sleep

Other than the role in ppGpp hydrolysis, Mesh1 has also been found to serve other functions in mammalian cells, such as NADPH phosphatase(Ding et al., 2020). To further investigate the role of ppGpp in sleep regulation, we ectopically expressed the *E. coli RelA* gene, which encodes a synthetase for ppGpp(Laffler and Gallant, 1974) in different set of neurons.

Firstly, we confirmed that the RelA from *E. coli* used by us indeed can increase ppGpp level *in vitro* (Fig. S1A). We then expressed RelA in neurons labeled by different drivers using Gal4-UAS system to specifically increase the ppGpp level in them. Because RelA and Mesh1 serve opposite functions in ppGpp metabolism, RelA expression would phenocopy *mesh1* mutant flies if mesh1 functions through ppGpp to regulate sleep. We found that when RelA was expressed in all the cells driven by *tub-Gal4* (O’Donnell et al., 1994), or in all the neurons driven by *elav-Gal4*(Robinow and White, 1991), sleep latency (Fig. 6A, columns 6 and 10) and awakening numbers at early night (Fig. 6B, columns 6 and 10) were significantly increased, phenocopying *M1KO* flies (Figs. 6A-B, column 2). By contrast, *UAS-RelA* driven by *repo-Gal4* for expression in glial cells(Halter et al., 1995) did not affect sleep (Figs. 6A-B, column 8). These results indicate that ppGpp functions in neurons, but not in glial cells, to regulate sleep.

**Figure 6.**
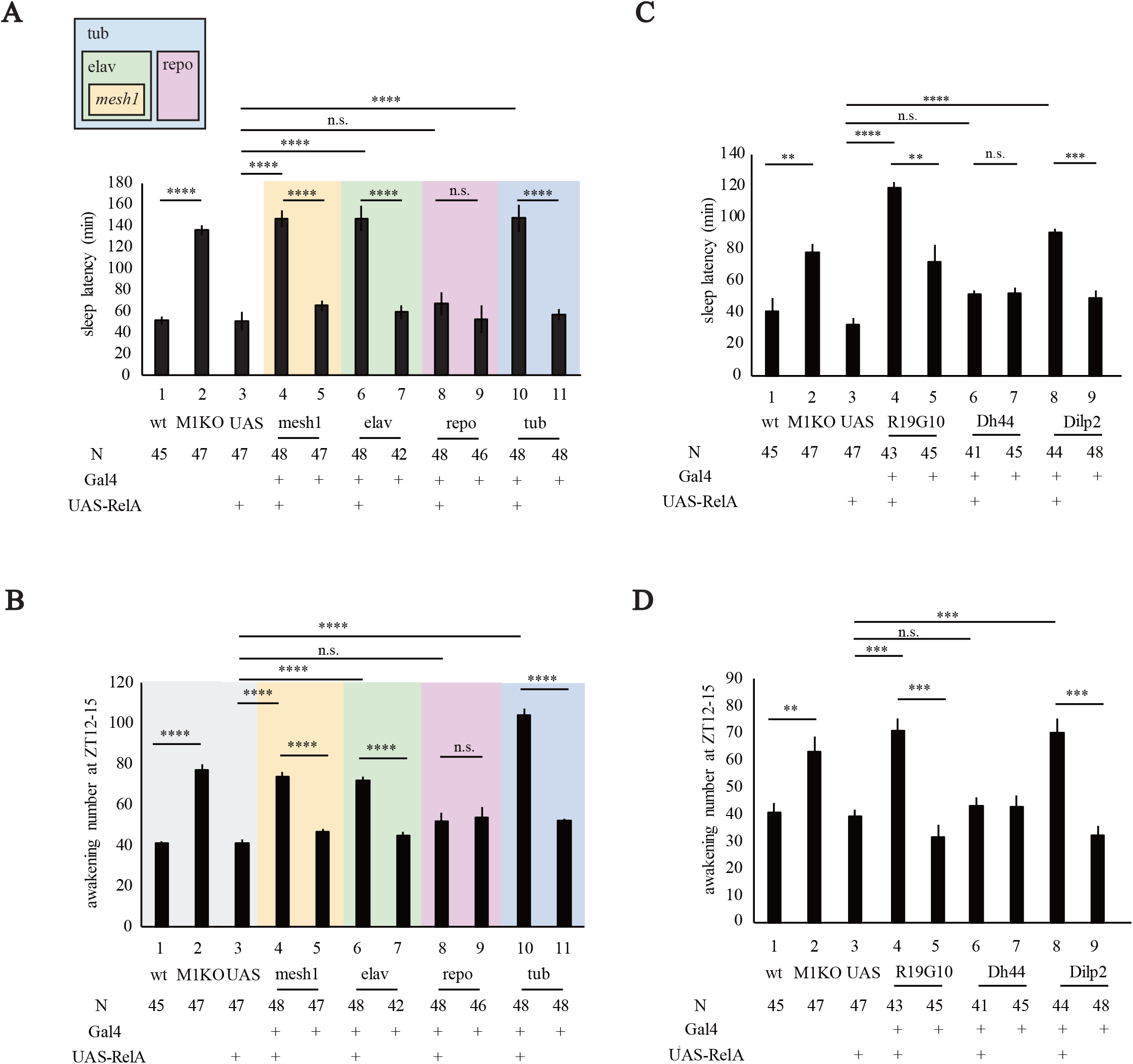
RelA ectopic expression in Dilp2 neurons phenocopies *mesh1* mutants in sleep. **(A-B)** Statistical analyses of sleep latency **(A)** and awakening numbers at early night **(B)** when RelA was expressed in all cells labeled by *tub-Gal4*, in neurons labeled by *elav-Gal4*, in glia labeled by *repo-Gal4*, or in *mesh1*-expressing cells labeled by *M1KOGal4*. Genotypes from left to right: **(1)** wildtype, **(2)** *M1KO/M1KO*, **(3)** *UAS-RelA/+*, **(4)** *M1KOGal4/UAS-RelA*, **(5)** *M1KOGal4/+*, **(6)** *elav-Gal4/+;;UAS-RelA/+*, **(7)** *elav-Gal4/+*, **(8)** *repo-Gal4/UAS-RelA*, **(9)** *repo-Gal4/+*, **(10)** *tub-Gal4/UAS-RelA*, **(11)** *tub-Gal4/+*. **(C-D)** Statistical analyses of sleep latency **(C)** and awakening numbers at early night **(D)** with RelA expression in different subsets of PI neurons. Genotypes from left to right: **(1)** wildtype, **(2)** *M1KO/M1KO*, **(3)** *UAS-RelA/+*, **(4)** *R19G10-Gal4/UAS-RelA*, **(5)** *R19G10-Gal4/+*, **(6)** *Dh44-Gal4/UAS-RelA*, **(7)** *Dh44-Gal4/+*, **(8)** *Dilp2-Gal4/+; UAS-RelA/+*, **(9)** *Dilp2-Gal4/+*. Two-way ANOVA and Bonferroni post-tests was used, ** *P* < 0.01, *** *P* < 0.001, **** *P* < 0.0001. Error bars represent s.e.m.

Furthermore, when RelA was specifically expressed in *mesh1*-expressing cells labeled by *M1KOGal4*, both sleep latency (Fig. 6A, column 4) and awakening numbers (Fig. 6B, column 4) were significantly increased, indicating that ppGpp level in *mesh1* positive neurons is sufficient to regulate sleep.

To test the effect of decreasing ppGpp level in different fly cells, we overexpressed *UAS-mesh1* in these drivers. Similar with RelA ectopic expression, *UAS-mesh1* overexpression driven by *tub-Gal4* in all cells or by *elav-Gal4* in neurons significantly decreased sleep latency (Fig. 7A, columns 5 and 9) and awakening numbers (Fig. 7B, columns 5 and 9), phenocopying flies with Mesh1 overexpressed in *mesh1*-expressing cells labeled by *Mesh1KOGal4* (Figs. 7A-B, columns 3). By contrast, *UAS-mesh1* overexpression driven by *repo-Gal4* in glial cells did not affect sleep latency or awakening numbers (Figs. 7A-B, column 7, Fig. 7C).

**Figure 7.**
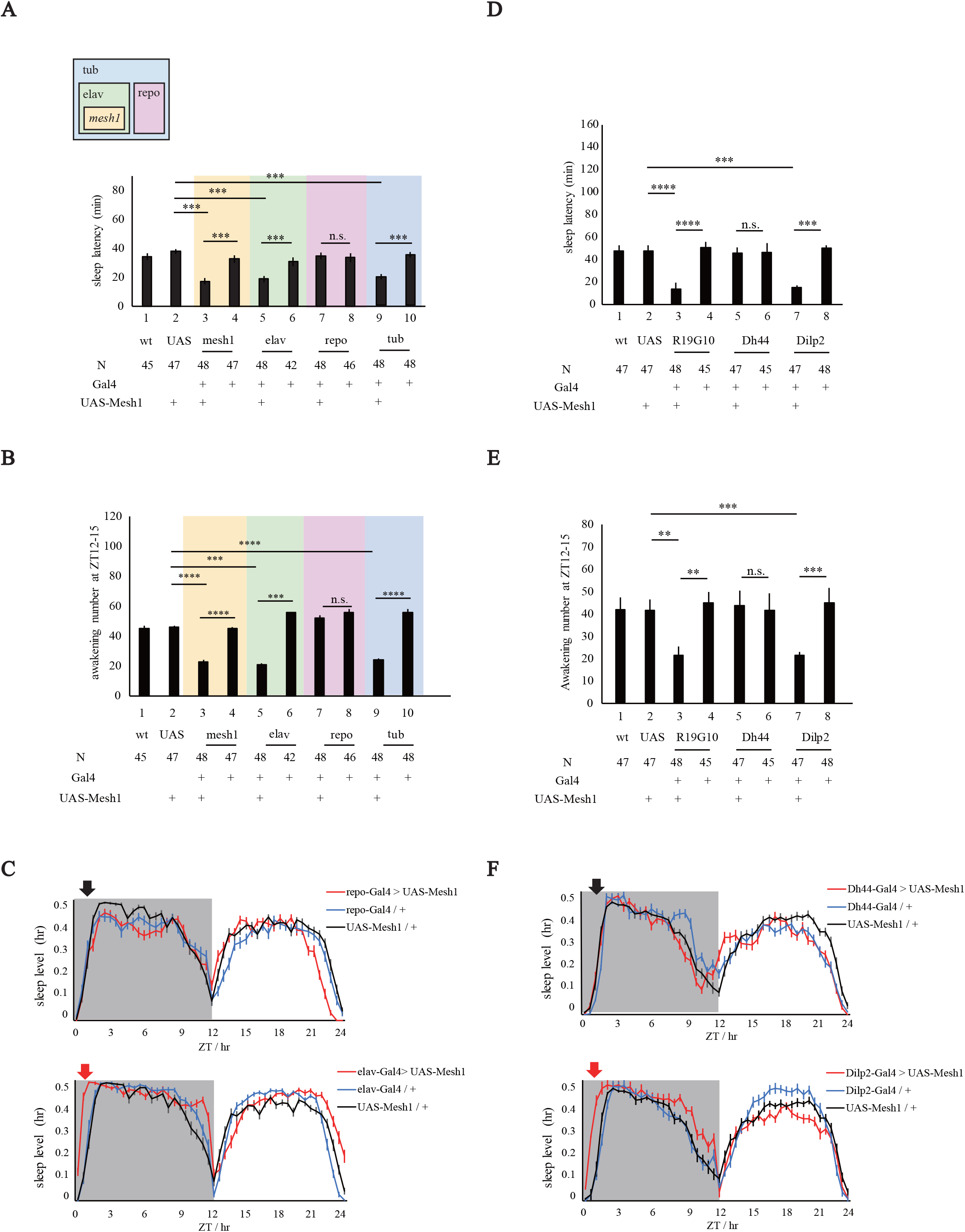
Mesh1 overexpression in subsets of neurons affected sleep. **(A-B)** Statistical analyses of sleep latency **(A)** and awakening numbers at early night **(B)** when Mesh1 was overexpressed in all cells, neurons, glia or *mesh1*-expressing cells. Genotypes from left to right: **(1)** wildtype, **(2)** *UAS-Mesh1/+*, **(3)** *M1KOGal4/UAS-Mesh1*, **(4)** *M1KOGal4/+*, **(5)** *elav-Gal4/+;;UAS-Mesh1/+*, **(6)** *elav-Gal4/+*, **(7)** *repo-Gal4/UAS-Mesh1*, **(8)** *repo-Gal4/+*, **(9)** *tub-Gal4/UAS-Mesh1*, **(10)** *tub-Gal4/+*. **(C)** Sleep profiles of lines in (A) and (B) in 30mins bins. Arrows denote sleep latency at early night. *UAS-Mesh1/+* in the top and bottom panels was the shared control, with the top panel showing that *repo-Gal4>UAS-Mesh1* did not affect sleep latency whereas the bottom panel showing that *elav-Gal4>UAS-Mesh1* shortened sleep latency. **(D-E)** Statistical analyses of sleep latency **(D)** and awakening numbers at early night **(E)** with Mesh1 overexpression in different subsets of PI neurons. Genotypes from left to right: **(1)** wildtype, **(2)** *UAS-Mesh1/+*, **(3)** *R19G10-Gal4/UAS-Mesh1*, **(4)** *R19G10-Gal4/+*, **(5)** *Dh44-Gal4/UAS-Mesh1*, **(6)** *Dh44-Gal4/+*, **(7)** *Dilp2-Gal4/+; UAS-Mesh1/+*, **(8)** *Dilp2-Gal4/+*. **(F)** Representative sleep profiles of lines in (D) and (E). Arrows denote sleep latency at early night. Note that the *UAS-Mesh1/+* profile in the top and bottom panels was the shared control, with the top panel showing that *Dh44-Gal4>UAS-Mesh1* did not affect sleep latency whereas the bottom panel showing that *Dilp2-Gal4>UAS-Mesh1* shortened sleep latency. Two-way ANOVA and Bonferroni post-tests was used, ** *P* < 0.01, *** *P* < 0.001, **** *P* < 0.0001. Error bars represent s.e.m.

Taken together, data from increasing or decreasing the ppGpp level by expressing RelA or Mesh1 with different drivers indicate that ppGpp functions in neurons but not in glia to regulate sleep.

### ppGpp functions in Dilp2 neurons to regulate sleep

Our results have shown that ppGpp functions in neurons, specifically, *mesh1*-expressing neurons, to regulate sleep (Figs. 4B-D, 6A-B, 7A-B). Next, to find where does ppGpp function to regulate sleep in flies, we made use of a Gal4 library of the chemoconnectome (CCT) previously generated by us, which include all the known neurotransmitters, modulators, neuropeptides and their receptors(Deng et al., 2019). We carried out a CCT screen by crossing each Gal4 line with *UAS-RelA*. We found that RelA expression driven by Gal4 lines of Trh, Capa-R, CCHa2-R, LkR, OA2 and CG13229 robustly affected sleep latency (Fig. S3A). It was noted that all of these lines drove expression in the PI region (Figs. S3B-G, insets), which suggests a functional significance of *mesh1* expression in the PI region(Fig. 5).

To further dissect the PI neurons for functional involvement in ppGpp regulation of sleep, we tested Gal4 lines known to drive expression in PI neurons: *Dh44-Gal4* (Cannell et al., 2016; Chen and Dahanukar, 2018), *Dilp2-Gal4* (Crocker et al., 2010; Semaniuk et al., 2018; Yurgel et al., 2019) and *R19G10-Gal4* (Collin et al., 2011; Ohno et al., 2017). To confirm whether *mesh1* is expressed in the PI neurons labeled by these drivers, we generated a Flp line of *mesh1* (*M1KIflp*, Fig. S4A), so that flippase will be expressed in *mesh1* cells and cut out the stop casette flanked by FRT in *UAS-FRT-stop-FRT-mCD8-GFP*. In this way, GFP is expressed in cells that are labeled by both the PI driver and *mesh1*. We found that mesh1 is indeed expressed in the PI neurons labeled by *Dh44-Gal4, Dilp2-Gal4*, and *R19G10-Gal4* (Figs. S4B-D). Also, co-expression of Dilp2 and Mesh1 was found by Dilp2 immunostaining of *M1KOGal4>UAS-mCD8-GFP* brains (Fig. S4E).

We then investigated the function of ppGpp in these PI neurons by crossing the drivers with *UAS-RelA* to increase ppGpp level in the neurons. Sleep latency (Fig. 6C) and awakening numbers (Fig. 6D) were significantly increased by RelA expression in neurons labeled by *R19G10-Gal4* and *Dilp2-Gal4*. However, RelA expression in Dh44 neurons did not affect sleep (Figs. 6C-D, columns 6).

Each of these PI drivers was crossed with *UAS-Mesh1* to decrease ppGpp level in the neurons. Consistent with previous results, sleep latency (Fig. 7D) and awakening numbers (Fig. 7E) were decreased by Mesh1 expression in neurons labeled by *R19G10-Gal4* and *Dilp2-Gal4*, but not in Dh44 neurons (Fig. 7F).

Taken together, experiments with RelA ectopic expression and Mesh1 overexpression have provided consistent results indicating that ppGpp functions in Dilp2 positive and Dh44-negative PI neurons to regulate sleep.

### ppGpp in Dilp2 neurons promotes starvation induced sleep loss

Sleep regulation gets input from many other activities, for example, starvation has been found to induce sleep loss in both humans(MacFadyen et al., 1973) and flies(Keene et al., 2010). Insulin signaling through Dilp2 plays a critical role in the interaction between diet and sleep homeostasis(Brown et al., 2020). Previous studies have suggested that ppGpp in bacteria and Mesh1 in flies is important to starvation response (Potrykus and Cashel, 2008; Ronneau and Hallez, 2019; Sun et al., 2010). Given the role of ppGpp in Dilp2 neurons in sleep regulation, we next investigated whether ppGpp regulates starvation induced sleep loss (SISL).

Baseline sleep in fed flies was recorded before flies were starved for 24 hours(Yurgel et al., 2019). Sleep during starvation was compared to that before starvation (Fig. 8A). Nighttime SISL in *M1KO* flies was significantly more than that in wt flies, whereas daytime SISL were similar between the wt and *M1KO* flies (Figs. 8A-B).

**Figure 8.**
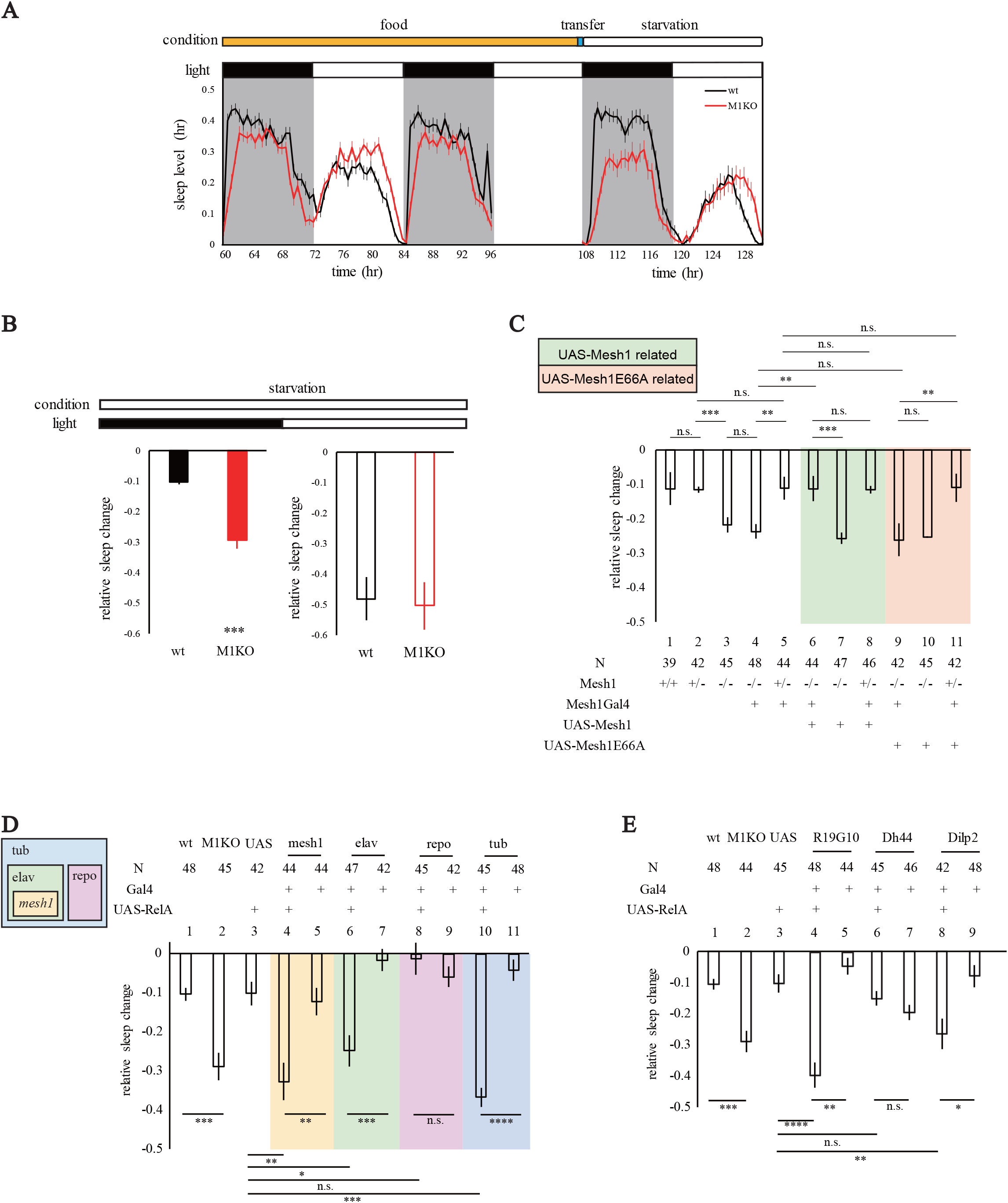
*mesh1* mutants and RelA ectopic expression in Dilp2 neurons both exacerbated SISL. **(A)** Sleep profiles of wt and *M1KO* flies before and after starvation. Flies were entrained for 3 days (recording at day 2 and day 3), transferred to 1% agar at the end of the 3rd day, followed by sleep recording with starvation of 24hrs. **(B)** Statistical analysis of SISL in (A). Sleep loss was significantly exacerbated in *M1KO* flies at nighttime. **(C)** Statistical analyses of SISL with Mesh1 and Mesh1E66A rescue experiments. Genotypes from left to right: **(1)** wildtype **(2)** *M1KO/+* **(3)** *M1KO/M1KO* **(4)** *M1KOGal4/M1KO* **(5)** *M1KOGal4/+* **(6)** *M1KOGal4/M1KO, UAS-Mesh1* **(7)** *M1KO/M1KO, UAS-Mesh1* **(8)** *M1KOGal4/UAS-Mesh1* **(9)** *M1KOGal4/M1KO, UAS-Mesh1E66A* **(10)** *M1KO/M1KO, UAS-Mesh1E66A* **(11)** *M1KOGal4/UAS-Mesh1E66A*. **(D)** Statistical analyses of SISL with RelA expression in all cells, neurons, glia or *mesh1*-expressing cells. Genotypes from left to right: **(1)** wildtype, **(2)** *M1KO/M1KO*, **(3)** *UAS-RelA/+*, **(4)** *M1KOGal4/UAS-RelA*, **(5)** *M1KOGal4/+*, **(6)** *elav-Gal4/+;;UAS-RelA/+*, **(7)** *elav-Gal4/+*, **(8)** *repo-Gal4/UAS-RelA*, **(9)** *repo-Gal4/+*, **(10)** *tub-Gal4/UAS-RelA*, **(11)** *tub-Gal4/+*. **(E)** Statistical analyses of SISL with RelA ectopic expression in different subsets of PI neurons. Genotypes from left to right: **(1)** wildtype, **(2)** *M1KO/M1KO*, **(3)** *UAS-RelA/+*, **(4)** *R19G10-Gal4/UAS-RelA*, **(5)** *R19G10-Gal4/+*, **(6)** *Dh44-Gal4/UAS-RelA*, **(7)** *Dh44-Gal4/+*, **(8)** *Dilp2-Gal4/+; UAS-RelA/+*, **(9)** *Dilp2-Gal4/+*. Student’s t-test was used in (B), two-way ANOVA and Bonferroni post-tests was used in (C)-(E), ** *P* < 0.01, *** *P* < 0.001, **** *P* < 0.0001. Error bars represent s.e.m.

We next investigated whether the enzymatic activity of Mesh1 is also important for SISL. The *M1KOGal4* mutants were similar to *M1KO* flies in having exacerbated SISL (Fig. 8C, columns 1-4 and Fig. S5A-C). *UAS-mesh1* (Fig. 8C, columns 6-8 and Fig. S5D) but not *UAS-mesh1E66A* (Fig. 8C, columns 9-11, and Fig. S5E) could rescue the phenotype of SISL in *mesh1* knockout flies, indicating that the ppGpp hydrolyzing activity is required for Mesh1 involvement in SISL.

Ectopic expression of the bacterial ppGpp synthetase RelA in Mesh1-expressing cells caused an exacerbated SISL (Fig. 8D, columns 4-5), phenocopying *M1KO* flies. Oppositely, expression of Mesh1 in Mesh1-expressing cells resulted in an alleviated SISL (Fig. 9A, columns 3-4). Taken together, these results indicate that ppGpp level, rather than any other unexpected activities of RelA or Mesh1 were involved in the SISL phenotypes.

**Figure 9.**
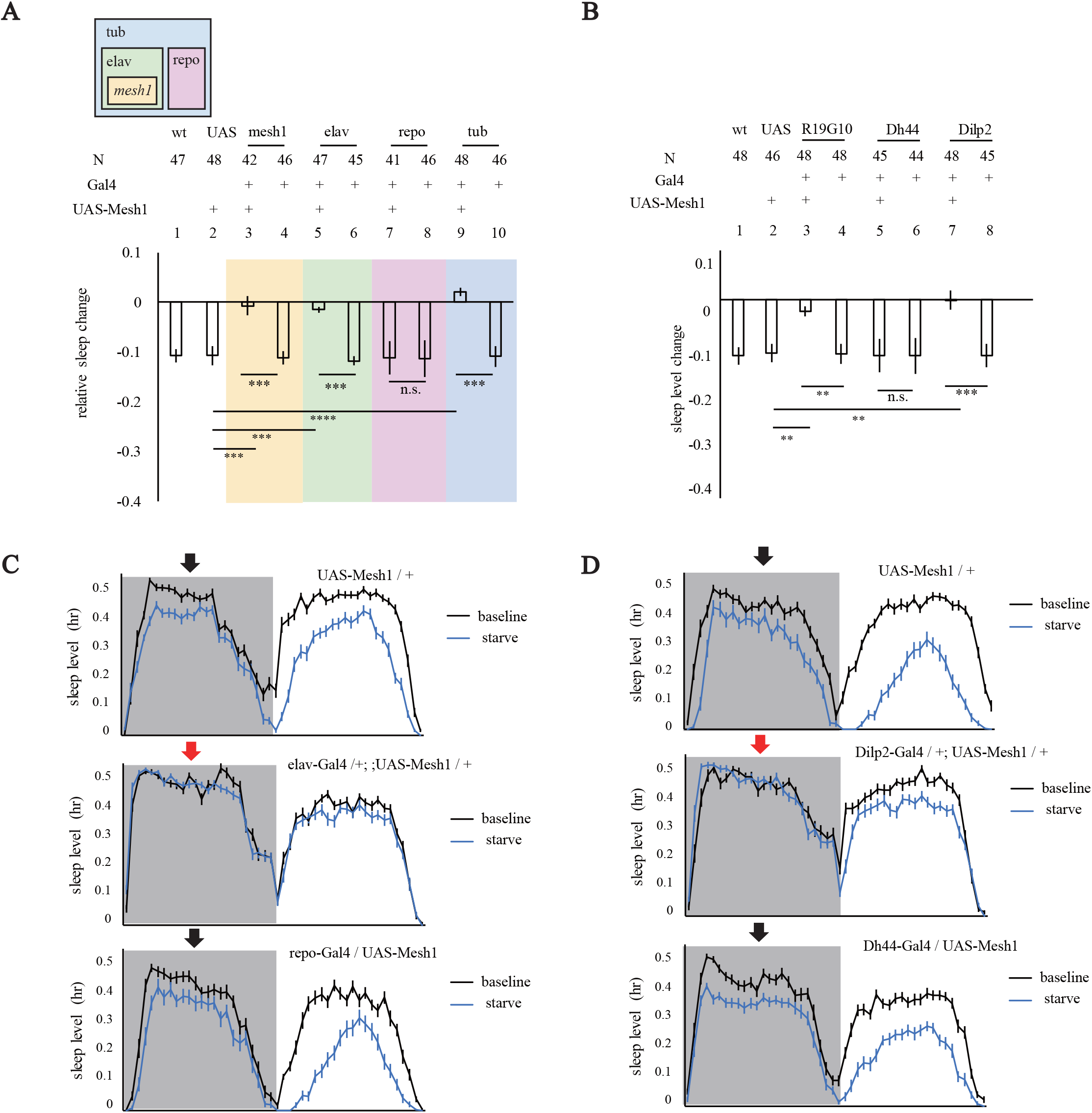
Mesh1 overexpression in Dilp2 neurons reduced SISL. **(A)** Statistical analyses of SISL with Mesh1 overexpression in neurons or other cells. Genotypes from left to right: **(1)** wildtype, **(2)** *UAS-Mesh1/+*, **(3)** *M1KOGal4/UAS-Mesh1*, **(4)** *M1KOGal4/+*, **(5)** *elav-Gal4/+;;UAS-Mesh1/+*, **(6)** *elav-Gal4/+*, **(7)** *repo-Gal4/UAS-Mesh1*, **(8)** *repo-Gal4/+*, **(9)** *tub-Gal4/UAS-Mesh1*, **(10)** *tub-Gal4/+*. **(B)** Statistical analyses of SISL with Mesh1 overexpression in different subsets of PI neurons. Genotypes from left to right: **(1)** wildtype, **(2)** *UAS-Mesh1/+*, **(3)** *R19G10-Gal4/UAS-Mesh1*, **(4)** *R19G10-Gal4/+*, **(5)** *Dh44-Gal4/UAS-Mesh1*, **(6)** *Dh44-Gal4/+*, **(7)** *Dilp2-Gal4/+; UAS-Mesh1/+*, **(8)** *Dilp2-Gal4/+*. **(C)** Representative sleep profiles of SISL in (A). Baseline sleep is in black, and sleep during starvation is in blue. Black arrows indicate that night SISL is comparable with wt; red arrows indicate night SISL was diminished. **(D)** Representative sleep profiles of SISL in (B). Two-way ANOVA and Bonferroni post-tests was used, ** *P* < 0.01, *** *P* < 0.001, **** *P* < 0.0001. Error bars represent s.e.m.

We then asked does ppGpp functions in the same set of cells to regulate both SISL and daily sleep. SISL was increased by general or neuronal expression, but not glial expression of RelA (Fig. 8D), and was decreased by general or neuronal overexpression, but not glial overexpression of Mesh1 (Figs. 9A and C), indicating that neuronal, but not glial, ppGpp regulates SISL. RelA (Fig. 8E) and Mesh1 (Figs. 9B) expression in PI neurons labeled by *Dilp2-Gal4* or *R19G10 Gal4* lines resulted in a change in SISL. By contrast, Dh44-expressing neurons were not involved in ppGpp regulation of SISL because neither RelA expression nor Mesh1 overexpression in Dh44 neurons affected SISL (Figure 8E, 9B and 9D).

Thus, the Dilp2 neurons that required for ppGpp regulation of sleep latency and awakening are also involved in its regulation of SISL.

## DISCUSSION

We have carried out a genetic screen of P element insertion lines which led to the discovery of a new mutation in the *Drosophila* ppGpp hydrolase Mesh1. Our chemical analysis revealed the presence of ppGpp in *Drosophila*. Our chemoconnectomic screen suggests involvement of PI neurons in ppGpp function. Further genetic intersection experiments confirm that ppGpp in specific PI neurons regulate sleep. Our findings indicate that ppGpp is present and has physiological functions in animals. It regulates sleep, including sleep latency and starvation induced sleep loss.

### Evidence for ppGpp Function in *Drosophila*

We obtained evidence for ppGpp function from four series of experiments. First, three mutations in mesh1 caused sleep phenotypes in Drosophila. The first mutation is a P-element insertion(Eddison et al., 2012), whose phenotype (Figs. 1A-B) and molecular nature we have characterized as *mesh1-ins* (Fig. 1C). The second mutation is a knockout generated by us as *M1KO* (Fig. 1C). The third mutation is a knockin generated by us as M1KOGal4 (Fig. 4A). All three lines have same phenotypes.

Second, we have shown that the hydrolyzing activity of Mesh1 is required to rescue the *mesh1* knockout mutant phenotype: if a point mutation was introduced to amino acid residue 66 by converting it from E to A, then it was enzymatically inactive *in vitro* (Fig. S1C) and unable to rescue the sleep and SISL phenotypes *in vivo* (Fig. 4, Fig. 8C).

Third, ectopic expression of RelA, a bacterial ppGpp synthetase, in *Drosophila* phenocopied *mesh1* knockout mutants (Fig. 6, Fig. 8).

Fourth, the phenotypes of Mesh1 overexpression are opposite to those of RelA expression (Fig. 7, Fig. 9).

### Sleep Regulation Role of ppGpp in Dilp2 Neurons

Does ppGpp function in all cells or only in some cells? Results obtained here support that ppGpp functions in specific neurons in the PI to regulate sleep.

*mesh1* gene is expressed in specific neurons (Fig. 5). Because Mesh1 is a ppGpp hydrolase, its expression can show the location where ppGpp is hydrolyzed, but not necessarily where it is synthesized or where it functions. When we screened Gal4 lines from CCT to drive the expression of RelA, sleep phenotypes were observed in 6 lines which were all expressed in the PI (Fig. S3). When Gal4 lines for PI neurons were used to drive RelA or Mesh1 expression, *Dilp2-Gal4* could indeed cause sleep and SISL phenotypes, while *DH44-Gal4* could not (Figs. 6-9). Expression of RelA or Mesh1 in glial cells did not affect sleep.

Because it is difficult to imagine that both synthetase and hydrolase could cause the same multiple phenotypes in the same neurons if these neurons are not where ppGpp functions, our results are most consistent with the idea that ppGpp functions in Dilp2 neurons to regulate sleep and SISL.

### ppGpp Synthesis in *Drosophila*

While we have shown that ppGpp is present in *Drosophila* and that it is hydrolyzed by Mesh1 which are expressed in specific neurons, we do not know how ppGpp is synthesized.

RSHs with the synthetase domain exist in both bacteria(Cochran and Byrne, 1974) and plants(van der Biezen et al., 2000), but not in animals(Atkinson et al., 2011; Sun et al., 2010). Either animals have evolved another enzyme to synthesize ppGpp, or that ppGpp in *Drosophila* is synthesized by bacteria and transported by specific mechanisms to neurons in the brain.

Plant RSHs are thought to result from lateral gene transfer events from bacteria(Field, 2018; Ito et al., 2017). Bioinformatic analyses and biochemical assays indicate that ppGpp synthetase homologs are distributed widely in plants including: dicotyledon *A. thalaniana*(van der Biezen et al., 2000), monocotyledon *O. sativa*(Xiong et al., 2001), green algae *C. reinhardtii*(Kasai et al., 2002), *S. japonica*(Yamada et al., 2003), *N. Tabacum*(Givens et al., 2004), pea plants(Takahashi et al., 2004), *P. nil*(Dabrowska et al., 2006), *C. annnum*(Kim et al., 2009), and moss *P. patens*(Sato et al., 2015). Presence of the N-terminal chloroplast transit peptide (cTP), causes most plant RSHs to be located in the chloroplasts(Boniecka et al., 2017; Chen et al., 2014; Mizusawa et al., 2008; Sato et al., 2009; Sugliani et al., 2016; Takahashi et al., 2004). It is unclear whether a ppGpp synthetase exists in *Drosophila* mitochondria.

### Molecular Targets of ppGpp

Previous studies have suggested a role of Dilp2 and Dilp2 neurons in sleep regulation under fed and starved conditions(Brown et al., 2020; Cong et al., 2015; Crocker et al., 2010). How does ppGpp function in these neurons to regulate sleep and SISL? In bacteria, the best-known direct target of ppGpp is RNA polymerase (RNAP)(Artsimovitch et al., 2004; Barker et al., 2001; Kajitani and Ishihama, 1984; Kingston et al., 1981; Lindahl et al., 1976). ppGpp interaction with the RNA polymerase leads to blockage of transcription initiation(Artsimovitch et al., 2004) and elongation(Kingston et al., 1981). There are also other possible targets for ppGpp(Corrigan et al., 2016; Dalebroux and Swanson, 2012; Gourse et al., 2018; Maciag et al., 2010; Nomura et al., 2014; Pao and Dyess, 1981; Paul et al., 2005; Sherlock et al., 2018; Zhang et al., 2019). A recent study utilizing a ppGpp-coupled bait uncovered new ppGpp target proteins in bacteria, including a large group of GTPase and metabolism-related enzymes(Wang et al., 2019). Future studies are required to identify the molecule(s) directly mediating sleep regulation of ppGpp in Dilp2 neurons in *Drosophila*.

## SUPPLEMENTAL INFORMATION

Supplemental information includes 5 figures and 1 table.

## ACKNOWLEDGEMENTS

We are grateful to Dr. U. Heberlein for sharing P-element insertion mutant library, Drs. J. Ni and G. Gao for providing us with CRISPR/Cas9 plasmids and flies, to Dr. Z.-F. Gong for sharing antibodies, to the Beijing Commission of Science and Technology and the National Natural Science Foundation of China (Project 31421003 to Y. R.) for grant support.

## AUTHOR CONTRIBUTIONS

Yi Rao supervised and initiated the project. Wei Yang performed the majority of experiments and data analysis. Xiaohui Zhang carried out experiments of UPLC-MS. Xihuimin Dai, Wei Yang, and Enxing Zhou carried out experiments of sleep screen on P-element insertion mutants. Wei Yang and Enxing Zhou implemented the fly tracing program and developed software for sleep analyses. Xihuimin Dai, Wei Yang and Yi Rao wrote the manuscript.

## METHODS

### Fly Stocks and Rearing Conditions

All flies were reared on standard corn meal at 25°C and 60% humidity, under 12hr:12hr LD cycle unless specified otherwise. Before behavioral assays, stocks were backcrossed into the background of an isogenized Canton S wt line in the lab for 7 generations.

Lines ordered from the Bloomington Stock Center included: #458 (elav-Gal4), #7415 (repo-Gal4), #47887 (R19G10-Gal4), #51987 (Dh44-Gal4), nos-phiC31, #37516 (Dilp2-Gal4), #5137 (UAS-mCD8-GFP). P-element insertion collection was a gift from Dr. U Heberlein (Janelia Research Campus)(Moore et al., 1998), and CCT-Gal4 library was a collection previously generated in our lab (Deng et al., 2019). isoCS and w^1118^ were wild-type and white-eye wild-type lines.

### Reagents and Plasmids

PCR was performed with Phanta-Max Super-Fidelity DNA Polymerase (Vazyme). Genotyping PCR was performed with 2x Taq PCR StarMix with Loading Dye (GenStar). Restriction enzymes KpnI-HF, SacII, NotI-HF, XbaI, XhoI, BamHI-HF, DpnI, EcoRI-HF and XbaI were from New England BioLabs. Total RNA was extracted from flies with RNAprep pure Tissue Kit (TIANGEN). Reverse transcription for cDNA cloning was performed with PrimeScript™ II 1^st^ Strand cDNA Synthesis kit (Takara), Gibson assembly was performed with NEBuilder HiFi DNA Assembly Master Mix. Transformation for cloning was performed with Trans-5 α (TransGen), and transformation for expression was performed with Transetta (TransGen). BL21 (TransGen) was used as the bacterial gene template. Reverse transcription for quantitative PCR (qPCR) analyses was performed with PrimeScript RT Master Mix kit (Takara). qPCR was performed with TransStart Top Green qPCR SuperMix kit (TransGen).

pACU2 was a gift from Drs. Lily and YN Jan. The plasmids previously used in our lab included: pBSK, pET28a+. Templates of STOP-attP-3Px3-RFP, T2A-Gal4-3Px3-RFP, and T2A-flp-3Px3-RFP were previously generated in the lab(Deng et al., 2019).

### Molecular Cloning and Generation of Transgenic Flies

Generation of all KO and KI lines was based on the CRISPR-Cas9 system with homologous recombination, according to previous procedures described(Ren et al., 2013). U6b vector was used for the transcription of sgRNA(Ren et al., 2013) and construction of targeting vectors were based on previous procedures in our lab(Deng et al., 2019).

To generate knock-out lines, a mixture of two *in vitro* transcribed U6b-sgRNA and one targeting vector was injected into *Drosophila* embryos. To generate U6b-sgRNA, two sgRNAs were selected on website (https://www.flyrnai.org/crispr), and designed into a pair of primers (M1KOSgRNA-1F, M1KOSgRNA-1R, M1KOSgRNA-2F and M1KOSgRNA-2R) without PAM sequence. The other pair of primers (U6b-laczrv and U6b-primer1) were designed to anneal the backbone of U6b vector, so that they could generate PCR products containing two parts of U6b (shorter fragment by M1KO sgRNA-F and U6b-laczrv, longer fragment by M1KO sgRNA-R and U6b-primer1), which share overlapping sequences in both ends. U6b-sgRNA plasmid was built by Gibson-assembly of the PCR products. To generate the targeting vector, two fragments (2kbps) flanking the entire CDS of gene Mesh1 except start codon were cloned as 5’Arm (amplified with genomic DNA with primers M1KO5F and M1KO5R) and 3’ arm (amplified from genomic DNA with primers M1KO3F and M1KO3R). Vector pBSK was digested with KpnI and SacII, and the PCR products were introduced into the digested pBSK by Gibson-assembly. New restriction sites (NotI and XhoI) were introduced between the two arms for further use. To generate the targeting vector for *M1KO*, STOP-attP-3Px3-RFP was amplified with primers attP2M1KOF and attP2M1KOR, and inserted between NotI and XhoI by Gibson-assembly; and for *M1KOGal4*, T2A-Gal4-3Px3-RFP was inserted at same location. A mixture of U6b-sgRNAs and the targeting vector was injected into embryos of *nano*-Cas9 or *vasa*-Cas9(Ren et al., 2013). F1 individuals of RFP+ eyes were collected after being crossed with w^1118^ flies.

To generate knock-in line *M1KIflp*, similar procedures were performed. For U6b-sgRNA, primers for shorter fragment were M1KISgRNA-1F or M1KISgRNA-2F paired with U6b-laczrv, while primers for longer fragment were M1KIsgRNA-1R or M1KIsgRNA-2R with U6b-primer1. For the targeting vector, 5’Arm was amplified with M1KI5F and M1KI5R, and 3’Arm with M1KI3F and M1KI3R. The site flanked by two arms was selected near the very end of Mesh1 CDS, so that the stop codon was removed and the CDS was fused with T2A-flp-3Px3-RFP with primers flp2M1KIF and flp2M1KIR.

Generation of transgenic UAS lines was based on vector pACU2(Han et al., 2011). *Drosophila* cDNA was generated by reverse transcription from total RNA. By using the cDNA as template, Mesh1 CDS was amplified with primers (M1CDSF and M1CDSR), and inserted into digested pBSK (by EcoRI and KpnI). Point mutation of Mesh1E66A was generated from this plasmid with primers M1E66AF and M1E66AR. Both Mesh1 CDS and Mesh1E66A were cloned with primers M1ACU2F and M1ACU2R. RelA sequence was cloned from BL21 bacteria with primers RAACU2F and RAACU2R. All the above were inserted into digested pACU2 (EcoRI with XbaI) by Gibson assembly. pACU2 constructs were inserted into attP2 by nos-phiC31 during embryo injection.

The seqence of primers are listed in Table S1.

### Molecular Cloning and Inducible Expression in Bacteria

pET28a+ was used for bacterial expression of Mesh1 and RelA proteins. Mesh1 CDS and Mesh1E66A were generated with primers M1ET28F and M1ET28R. RelA was cloned from BL21 bacteria with primers RAET28F and RAET28R. They were inserted into pET28a+ by Gibson assembly. Competent cells were transformed with the above constructs, and the strains were grown for inducible expression. To induce expression, a colony was inoculated into 1ml kanamycin containing Luria broth (LB) medium, and incubated at 37°C for 2 hr. 800μl was transferred into a flask with 800ml kanamycin+ LB media and incubated for ∼3hr at 37°C 220rpm, until OD_600_ was 0.5 to 0.6. 800μl 1M isopropyl β-D-1 thiogalactopyranoside (IPTG) was added into the flask. For RelA, the induction was at 37°C for 3hr; for Mesh1 and Mesh1E66A, it was at 16°C for 16 hr. Bacteria was harvested by centrifugation followed by lysis with ultrasonication (Power: 30%, lysis 2s, wait 2s, 30 cycles) and centrifugation. Supernatants were passed through nickel columns, followed by two times of wash with 10ml binding buffer (20mM Tris-HCl pH 7.4, 0.5M NaCl, 5mM imidazole), and eluted with 5ml elution buffer (20mM Tris-HCl pH7.4, 0.5M NaCl, 500mM imidazole). Each step was monitored on 10% SDS-PAGE to check protein expression. Eluted protein was enriched in Millipore Amicon Ultra-15, and resuspended in 1ml of protein storage buffer (20mM Tris-HCl pH7.4, 150mM NaCl, 0.3% CHAPS, 1mM DTT).

### Extraction and Measurement of ppGpp

To extract ppGpp from Drosophila, 3ml formic acid was added to every 250 flies. After grinding for 15 seconds, 1ml 30% tri-chloric acetic acid was added to precipitate proteins. The above procedures were repeated to collect extracts from 2000 flies. Supernatants were lyophilized overnight, and powders were resuspended with 200μl water.

To measure the level of ppGpp, C18 column was used in UPLC-MS, according to a modified version of previous method(Ihara et al., 2015). The mobile phase included: Buffer A (8mM *N,N*-dimethylhexylamine, 160μl acetic acid, 500ml water) and Buffer B (Acetonitrile). The UPLC program was: at 0min, A:B = 100% : 0%; at 10min, A:B = 40% : 60%, with linear increment. The m/z of ppGpp should be 601.95, and ATP (506.00) was used for normalization.

### Behavioral Assays

To analyze baseline sleep under 12hr:12hr L:D cycles, approximately 48 flies of each genotype were loaded into glass tubes for video tracing (fps=1), which was analyzed by an in-house software as described previously(Dai et al., 2019; Deng et al., 2019; Qian et al., 2017). Continuous immobility of >5min was defined as a sleep bout(Hendricks et al., 2000; Shaw and Brody, 2000). Sleep latency at early night was defined as the time from light-off (ZT12) to the point when the 1^st^ sleep bout appeared.

To analyze awakening numbers, video traced data was converted to data of simulated beam-crossing. In the simulation, the middle line for each tube was set as the virtual beam. According to a previous study(Tabuchi et al., 2018), brief awakening was defined as 1 cross per min, and the awakening number was the sum of such events in every 30 min. Awakening number of early night was the sum of brief awakenings at ZT12-15, and awakening number of late night was the sum of brief awakenings at ZT21-24.

To test starvation-induced sleep loss (SISL), sleep was recorded for the first 3 days after flies were loaded into tubes(the 3^rd^ day defined as baseline), and then flies were quickly transferred to tubes of 1% agar at the end of the light phase of 3^rd^ day, followed by a 24 hr recording of sleep during starvation. SISL ratio was defined as (starvation sleep-baseline sleep)/ baseline sleep.

### Immunohistochemistry and Imaging

To prepare flies for imaging, 5 flies per genotype were dissected in phosphate-buffered saline (PBS). Dissected tissues were transferred to a tube of 400μl 2% PFA, and fixed for 55 min. Tissues were washed 3 times with 400μl brain wash buffer (PBS containing 1% TritonX-100, 3% m/V NaCl), and then transferred to 400μl blocking buffer (PBS containing 2% TritonX-100, 10% normal goat serum), followed by incubation at 4°C overnight. Tissues were then transferred to dilution buffer (0.25% TritonX-100, 1% NGS, 1x PBS) and added with primary antibodies. Tissues were stained at 4°C overnight, followed by 3 washes with 400μl brain wash buffer. Samples were transferred to fresh dilution buffer containing secondary antibodies (1:200 Alexa Fluor goat anti-chick 488 (Invitrogen) and 1:200 Alexa Fluor goat anti-mouse 633 (Invitrogen)), followed by 3 washes with 400μl brain wash buffer. Samples were then mounted on slides in Focus Clear (Cell Explorer Labs, FC-101), and imaged on Zeiss LSM 710 confocal microscope.

Chicken anti-GFP (1:1000) (Abcam Cat# 13970; RRID:AB_300798) and mouse anti-Bruchpilot (1:40) (DSHB Cat# 2314866, nc82; RRID: AB_2314866) were used as primary antibodies with AlexaFluor488 anti-chicken (1:500) (Life Technologies Cat# A11039; RRID:AB_2534096) and AlexaFluor633 anti-mouse (1:500) (Life Technologies Cat# A21052; RRID: AB_141459) being used as respective secondary antibodies.

## CONTACT FOR REAGENT AND RESOURCE SHARING

Further information and requests for resources and reagents should be directed to and will be fulfilled by the Lead Contact, Yi Rao.

Strains and plasmids are available upon request. The authors affirm that all data necessary for confirming the conclusions of the article are present within the article, figures, and tables.

## FIGURE LEGENDS

**Figure S1. ppGpp quantification and validation of germ-free flies**.

**(A)** ppGpp level *in vitro*. From left to right: **(1)** Standard ppGpp, a commercial sample;

**(2)** Substrate only, the mixture of reaction buffer, GDP and ATP; **(3)** RelA+GDP, purified RelA was added to the substrate; **(4)** Mesh1+ppGpp, standard ppGpp and purified Mesh1 were added to the substrate; **(5)** Mesh1E66A+ppGpp, standard ppGpp and purified Mesh1E66A were added to the substrate.

**(B)** Schematic representations of the ppGpp synthetase domain (SD) and hydrolase domain (HD) in RSH proteins from *E. coli, D. melanogaster*, and *H. sapiens*. The mammalian Hddc3 and the *Drosophila* Mesh1 proteins contain only HD, the bacterial RelA protein contains a weak HD, an active SD and a regulatory domain (RD). Bacterial SpoT contains an active HD, a weak SD and a RD.

**(C)** Examination of germ-free status by growing bacteria on LB plate. 6 conditions were examined by spread LB plate with: **(1)** blank: nothing; **(2)** water germ-: germ-free water; **(3)** wt germ-: extract of germ-free wt flies; **(4)** M1KO germ-: extract of germ-free *M1KO* flies; **(5)** wt germ+: extract of normal wt flies; **(6)** M1KO germ+: extract of normal *M1KO* flies. After incubation, bacteria were only found on plates of wt germ+ and M1KO germ+ among these conditions.

**(D)** Examination of germ-free status by 16S rDNA PCR. 5 pairs of 16S PCR primers were used, including **(1)** 16S-27F/16S-519R; **(2)** 16S-357F/16S-907R; **(3)** 16S-530F/16S-1110R; **(4)** 16S-926F/16S-1492R; **(5)** 16S-1114F/16S-1525R. The upper and lower bands at ladder lane (Trans2K) on the left correspond to 500bp and 250bp. PCR signals by primers 1, 3, and 5 were detected in wt germ+ and M1KO germ+, while no signals were detected in all three germ-free groups (water germ-, wt germ-, M1KO germ-).

**(E)** ppGpp level in flies. From left to right, the genotypes are: **(1)** *Mesh1KO/+* **(2)** *Mesh1KO/Mesh1KO* **(3)** *Mesh1KOGal4/+* **(4)** *Mesh1KOGal4/ Mesh1KO, UAS-Mesh1* **(5)** *Mesh1KOGal4/UAS-Mesh1* **(6)** *Mesh1KOGal4/ Mesh1KO, UAS-Mesh1E66A* **(7)** *Mesh1KOGal4/UAS-Mesh1E66A*

**Figure S2. Sleep bout number and length of *M1KO* flies**

Statistical analyses of sleep bout number **(A-B)** and bout length **(C-D)** in nighttime **(A, C)** and daytime **(B, D)**. Sleep bout number and bout length of *M1KO* flies were comparable with wt flies. Student’s t-test was used. Error bars represent s.e.m.

**Figure S3. A screen of RelA ectopic expression in CCT-Gal4 lines**

**(A)** The result of screen for sleep latency with 102 CCT-Gal4 lines driving RelA expression. Candidates above blue line showing three standard deviations away from the mean. RelA expression in cells labeled by 6 Gal4 lines were found to significantly increase sleep latency: Capa-R, CCHa2-R, LkR, OA2, CG13229 and Trh.

**(B-G)** mCD8-GFP expression driven by each of the CCT Gal4 lines: Capa-R **(B)**, OA2 **(C)**, CCHa2-R **(D)**, CG13229 **(E)**, LkR **(F)** and Trh **(G)**, immunostained with anti-GFP (green) and nc82 (magenta). Insets: PI neurons labeled by each line.

**Figure S4. Mesh1 expression in PI neurons**

**(A)** Schematic representation of *M1KIflp*. 2A-flp-3Px3-RFP was fused in-frame to the C terminus of *mesh1*.

**(B-D)** Expression patterns of *M1KIflp, UAS-FRT-STOP-FRT-mCD8-GFP* in the brain, immunostained with anti-GFP (green) and nc82 (magenta). Cells expressing mesh1 and the PI-Gal4 were labeled by GFP. Insets: higher magnification views of PI.

**(E)** Co-expression of Dilp2 and Mesh1. *M1KOGal4>UAS-mCD8-GFP* brains were immunostained with anti-GFP (green) and anti-Dilp2 (red).

**Figure S5. Sleep profiles of SISL rescue experiments**

Baseline sleep is shown in black, and sleep during starvation shown in blue; black arrows indicate that night SISL is not significantly changed compared to wt or parental controls; red arrows indicate that a specific genotype phenocopied M1KO in SISL. Exacerbated SISL in mesh1 mutants **(A-C)** were rescued by Mesh1 expression in mesh1 cells **(D)**, but could not be rescued by Mesh1E66A expression **(E)**.

**Table S1. A list of used primers**.

